# Effect of Tokenization on Transformers for Biological Sequences

**DOI:** 10.1101/2023.08.15.553415

**Authors:** Edo Dotan, Gal Jaschek, Tal Pupko, Yonatan Belinkov

## Abstract

Deep learning models are transforming biological research. Many bioinformatics and comparative genomics algorithms analyze genomic data, either DNA or protein sequences. Examples include sequence alignments, phylogenetic tree inference and automatic classification of protein functions. Among these deep learning algorithms, models for processing natural languages, developed in the natural language processing (NLP) community, were recently applied to biological sequences. However, biological sequences are different than natural languages, such as English, and French, in which segmentation of the text to separate words is relatively straightforward. Moreover, biological sequences are characterized by extremely long sentences, which hamper their processing by current machine-learning models, notably the transformer architecture. In NLP, one of the first processing steps is to transform the raw text to a list of tokens. Deep-learning applications to biological sequence data mostly segment proteins and DNA to single characters. In this work, we study the effect of alternative tokenization algorithms on eight different tasks in biology, from predicting the function of proteins and their stability, through nucleotide sequence alignment, to classifying proteins to specific families. We demonstrate that applying alternative tokenization algorithms can increase accuracy and at the same time, substantially reduce the input length compared to the trivial tokenizer in which each character is a token. Furthermore, applying these tokenization algorithms allows interpreting trained models, taking into account dependencies among positions. Finally, we trained these tokenizers on a large dataset of protein sequences containing more than 400 billion amino acids, which resulted in over a three-fold decrease in the number of tokens. We then tested these tokenizers trained on large-scale data on the above specific tasks and showed that for some tasks it is highly beneficial to train database-specific tokenizers. Our study suggests that tokenizers are likely to be a critical component in future deep-network analysis of biological sequence data.

## Introduction

Since the development of modern DNA sequencing technologies, there has been a rapid growth of available genomic data. While a relatively small bacterial genome such as *Escherichia coli* is roughly five million bases (Markowitz et al. 2012), the complete sequence of a human genome is more than three billion bases long (Nurk et al. 2022). Current large-scale protein datasets are growing at an exponential rate and already encompass hundreds of billions of amino acids (Steinegger and Söding 2018). In light of the increasing size and length of biological sequence datasets, new processing methods are needed.

Deep-learning algorithms transformed many fields (LeCun, Bengio, and Hinton 2015), including computer vision (Voulodimos et al. 2018), biomedical research (Rudas et al. 2023), and comparative genomics (Miller, Stern, and Burstein 2022). The use of deep learning for genomic analysis is a game-changer and gains momentum as neural network solutions usually outperform traditional algorithms (Jumper et al. 2021; Kulmanov, Khan, and Hoehndorf 2018). Both natural human languages and biological sequences are composed of discrete characters (letters and nucleotides, respectively). These characters are the building blocks of sophisticated structures, i.e., text and genomes, which include elements such as sentences and genes, respectively. Although NLP architectures can be adapted to biological problems, considerable differences remain between human language and genomic data (Yu et al. 2019; List et al. 2016; Dotan et al. 2023). Among the major differences are the sequence length and the size of the dictionary, i.e., the entire set of tokens used in that language. Human languages usually contain short sequences with a large dictionary, e.g., a dictionary size of hundreds of thousands of words in English. In contrast, genomic data contain considerably longer sequences with a much smaller dictionary, a dictionary size of four for the DNA sequences.

Long sequences raise memory consumption and run-time challenges when analyzed using deep-learning networks. Different approaches to tackle these issues have emerged, including: (1) Developing specific architectures for long sequences (Lin et al. 2021; Rao et al. 2021); (2) Splitting the data into smaller segments (Dotan et al. 2023); (3) *K*-mer representation of all possible nucleotides (Ji et al. 2021).

Deep-learning architectures for traditional NLP tasks frequently operate with a fixed vocabulary and thus have a limited size. As the number of different words in human languages often exceeds this fixed size, some of the words may be considered unknown. Tokenizers for human languages emerged as a solution to reduce the number of unknown words, i.e., words that are not part of the vocabulary. For example, the word “Bioinformatics”, might be split into three subwords, “Bioin”, “form”, and “atics”. The tokenization process is data driven and hence does not necessarily correspond to linguistically meaningful units. Tokenization algorithms split such words into common subwords and thus enable NLP-based methods to put these tokens in context. This may result in increased ability to infer meanings. Subwords tokenizers include “Byte-Pair Encoding” (BPE), “WordPiece” and “Unigram” (Sennrich, Haddow, and Birch 2016; Schuster and Nakajima 2012; Kudo 2018). The BPE and WordPiece tokenizers initialize a dictionary consisting of all the characters in the raw text, and progressively select pairs of tokens to merge and add them as a new token to the dictionary. BPE and WordPiece differ in how pairs are selected: while BPE adds the most frequent pair, WordPiece adds the pair that maximizes the frequency of the pair divided by the product of the frequencies of the two tokens (see Methods). The Unigram methodology is different. It initializes a dictionary consisting of a very large number of relevant tokens. The dictionary is next trimmed by removing non-contributing tokens, which are inferred by applying a specific loss function (see Methods).

As mentioned above, biological data behave differently as the base “words” are often nucleotides or amino acids and hence cannot be broken any further. This is contrary to human languages, in which some tokenizers break the words into characters. For example, when using the dictionary of the four nucleotides, “A”, “C”, “G”, and “T”, the sequence length of an entire genome will be the number of nucleotides (Figure 1a). One can think of different dictionaries, e.g., one that contains all the possible pairs: “AA”, “AC”, “AG”, “AT”, “CA” … “TT”. This raises the size of the dictionary by a power of two (i.e., 16) and reduces the length of the sequences by approximately two folds (Figure 1b). Of note, the conversion of nucleotides to codons has a similar impact as the dictionary size is 64 (61 sense codons and three stop codons) and the sequence length is reduced by a factor of three.

**Figure 1:**
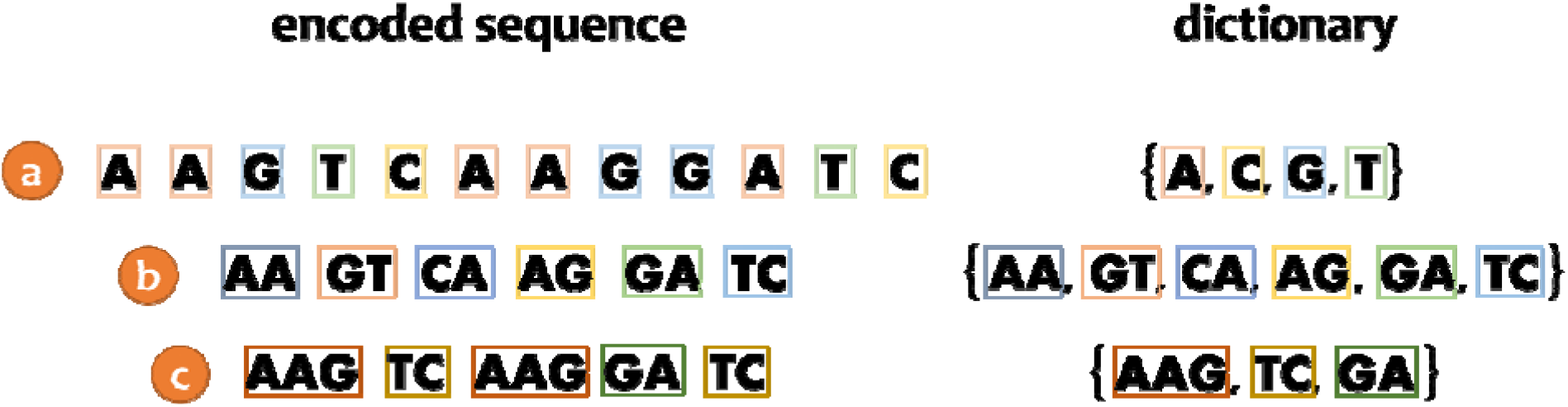
Different tokenization algorithms can be applied to biological sequences, as exemplified for the sequence “AAGTCAAGGATC”. (a) The baseline “words” tokenizer assumes a dictionary consisting of the nucleotides: “A”, “C”, “G” and “T”. The length of the encoded sequence is 12, i.e., the number of nucleotides; (b) The “pairs” tokenizer assumes a dictionary consisting of all possible nucleotide pairs. The length of the encoded sequences is typically halved; (c) A sophisticated dictionary consisting of only three tokens: “AAG”, “TC” and “GA”. The encoded sequence for this dictionary contains only five tokens.

The genome of each species contains unique substrings, as well as various repetitive elements. These repetitive elements, often named *K*-mers, vary in type and length. We expect data-driven biological tokenizers to assign a token for such repetitive elements. Of note, repetitive elements comprise more than half of the human genome (Richard, Kerrest, and Dujon 2008). The existence of such *K*-mers across multiple species motivates the employment of data driven tokenizers which can substantially reduce the number of tokens to process without a substantial increase in the size of the dictionary. Of note, reducing the number of tokens partially alleviates the problems encountered with long sequences. However, too large dictionaries forbid capturing shared elements among sequences. Which tokenizer best balances between these two constraints is an open question. In this study, we focus on comparing transformers trained on data processed by different tokenizers, in terms of performance and input length.

## Methods

### Outline

We aim to train and evaluate alternative tokenizers. The input is a set of biological sequences. Different biological tasks are considered, e.g., classifying the sequence into several categories. The tokenizers are applied to the input sequences creating a list of integers, which represent the different tokens. Such lists are used to train a deep-learning network model (in our case, a transformer). A single transformer (Vaswani et al. 2017) is trained for each biological task and each tokenizer. The performance is measured on transformer processing test data (which were processed with the same tokenizer as the training data). In addition, we report the effect on the number of input tokens, which is a proxy for memory and runtime consumption.

### Tokenizers

We evaluated five different tokenizers on biological sequences: BPE, Unigram, WordPiece, “words”, and “pairs”. “Words” is a trivial tokenizer, in which the dictionary contains all possible amino-acids or nucleotides. In the “pairs” tokenizer, the dictionary contains all possible pairs of characters (amino-acids or nucleotides). Of note, while in “words” and “pairs” the dictionary size is fixed, in the other three tokenizers, the dictionary size is a tunable parameter of the model. Thus, we tested different values of dictionary size. For each computational task and for each combination of tokenizer and dictionary size, a different transformer was trained, and its performance evaluated (described below). We would like to emphasize the differences in applying tokenizers to natural languages versus biological sequences. In most natural languages there are three levels of text representation: characters, words, and sentences. In contrast, in biological sequences, there are only two levels, as they lack the space character separating words in natural languages.

#### BPE (Byte-Pair Encoding)

Initially, BPE was a general purpose compression algorithm (Gage, 1994), but it has since been adopted for tokenizing textual data, initially for machine translation with recurrent neural networks (Sennrich, Haddow, and Birch 2016) and later for transformers (Vaswani et al. 2017). The tokenizer creates a base vocabulary from unique characters in the pre-tokenized data, and then gradually merges the most frequent pair, adding each new one to the vocabulary. This process stops when the vocabulary size reaches a hyperparameter that must be determined before training the tokenizer.

#### WordPiece

This tokenizer is similar to BPE (Schuster and Nakajima 2012). Like BPE, it uses the entire set of characters in pre-tokenized data to create a base vocabulary and progressively adds new tokens to it. Unlike BPE, which adds the most frequent pair, WordPiece selects the pair that maximizes a certain score:

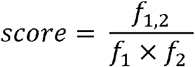

*f*_1,2_ is the frequency of the pair of elements, while *f*_1_ and *f*_2_ are the frequencies of the two separate elements.

#### Unigram

Unigram (Kudo 2018) takes a different approach than BPE and WordPiece. It begins with a heuristic identification of an initialized vocabulary, which is later trimmed. There are different ways to create the initial vocabulary, e.g., selecting the most frequent sub-strings in the corpus or using a different tokenizer such as BPE with specific hyperparameters that yield a large vocabulary. Next, the Unigram tokenizer progressively removes tokens from the vocabulary by searching for tokens whose removal improves the model fit, as quantified using a loss function (detailed below). Usually, more than one token is removed at a time, since computing the loss for all tokens is a costly operation. Given a corpus of *N* words, *x*_1_, …, *x*_*i*_, …, *x*_*N*_, the loss is the sum of the negative log-likelihood of the score of each word, denotated by *h*(*x*_*i*_) for the word *i*:

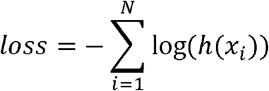

where *h*(*x*_*i*_) is the maximum score of dividing the word *x*_*i*_ to tokens:

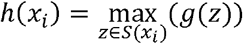

*S*(*x*_*i*_) are all the possible options to split *x*_*i*_ to tokens, and *g* maps a specific set of tokens, i.e., option, *t*_1_, …, *t*_*j*_ …, *t*_*M*_ = *z* to a score.

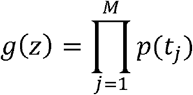

Where *p*(*t*_*j*_) is the unigram probability of token *j*, i.e., the number of occurrences of token *j* divided by the total number of tokens in the corpus.

### Biological Datasets

We compared the performance of the above tokenizers on eight datasets, described below, which vary in terms of their size, the learning task required (five classifications, two regressions, and one sequence alignment), and the type of sequence data (one dataset contains DNA sequences, while the others contain protein sequences).

#### Dataset1

Type III effectors (we will use the term effectors below) are proteins that are secreted by pathogenic bacteria from the bacterial cytoplasm into the host cell. In the host cell, they manipulate cellular processes to the benefit of the bacteria. The computational challenge is to classify bacterial proteins to those that are effectors and those that are not. The secretion signal that determines whether a protein is an effector or not was shown to reside in the 100 amino acids of the N-terminus of a protein (Notti and Stebbins 2016). We obtained a dataset of 641 effectors and 4,544 non-effector proteins (Wagner et al. 2022). From each protein, we only considered the 100 N-terminal amino acids. The true label (whether or not the protein is an effector) is known from experimental work. The computational task is to correctly classify each protein to its category. These data were divided to training, validation and test, each containing 497, 60 and 84 effectors and 2,034, 219 and 2,291 non-effectors, respectively.

#### Dataset2

A superfamily is a group of proteins that share similar properties and functions. The second task is to classify proteins to superfamilies based on their amino-acid sequences. To this end, we downloaded sequences from the Pfam database (Mistry et al. 2021). We randomly picked nine different superfamilies containing over 2,000 protein sequences: (1) SSF100895 Kazal-type serine protease inhibitors; (2) SSF110035 GDNF receptor-like; (3) SSF109993 VPS9 domain; (4) SSF101152 Mob1/ phocein; (5) SSF110019 ERO1-like; (6) SSF102546 RbsD-like; (7) SSF101912 Sema domain; (8) SSF100939 SPOC domain-like; (9) SSF100879 Lesion bypass DNA polymerase (Y-family), little finger domain. For each superfamily we downloaded the first 2,000 protein sequences, which we split into training, validation and test data, containing 1,800, 100, and 100 sequences, respectively. The Pfam database includes information regarding the specific fragments issued with the superfamily. For each of the sequences, we extracted these fragments and concatenated them. The task is predicting the superfamily given the fragments.

#### Dataset3

Sequence alignment is one of the common tasks in bioinformatics (Van Noorden, Maher, and Nuzzo 2014) as it provides a record of similarity between homologous sequences. One has to account for different evolutionary events such as insertions, deletions and substitutions to correctly infer the alignment. The third dataset contains pairwise homologous nucleotide sequences, and the task is to correctly align them. We have previously developed a deep-learning-based algorithm for such an alignment task, in which we train transformers to map pairs of unaligned sequences, i.e., source sentences, into a valid alignment, i.e., target sentences (Dotan et al. 2023). The average number of nucleotides is 429 and 434 for the source and target sentences, respectively. This dataset was simulated by SpartaABC (Loewenthal et al. 2021), and hence the correct alignment is known. The data include 395,000, 2,000 and 3,000 training, validation and test alignments, respectively. To simulate those sequences, we have used the following parameters: (1) Root length between 150 to 300 nucleotides; (2) Pairwise evolutionary distance between 0.05 to 0.15 substitutions per site (3) An insertion rate between 0.0 to 0.05 events per substitution and similarly for deletions; (4) The “A parameter” dictates the length distribution of insertion and deletion events. The A parameter for insertions ranged between 1.01 and 2.0, and similarly for deletions. For each simulation of a pair of sequences, SpartaABC samples uniformly from those ranges and generates the alignment based on the sampled parameters.

#### Dataset4

Protein folds are characteristics of the protein three-dimensional structure. Often, proteins evolve so that their sequence similarity becomes low, yet, they still share substantial structural similarity. Nevertheless, the folding information is encoded within the protein sequence. Here we analyzed 13,766 protein sequences, each of which is labeled by a specific fold (Hou, Adhikari, and Cheng 2018). There is a total of 1,195 protein folds, and the computational task it to classify each protein to its correct fold based on its amino-acid sequence. These data were partitioned to 12,312, 736, 718 training, validation, and test pairs of sequence-fold.

#### Dataset5

The fluorescence intensity of a protein is determined by its sequence and structure. The general mapping form sequence space to fluorescence intensity is unknown in general. Here we rely on previously established data (Sarkisyan et al. 2016), which were partitioned to 21,446, 5,363, and 27,217 training, validation, and test pairs of sequence-log-intensity values, respectively. Of note, unlike the previous tasks, here a regression model is needed from the sequence space to fluorescence intensities.

#### Dataset6

In stability landscape prediction, the challenge is to predict the concentration threshold from which the protein unfold, given the sequence of amino-acids (Rocklin et al. 2017). For this regression task, the data included 53,614, 2,512, 12,851 training, validation, and test pairs of sequence-concentration, respectively.

#### Dataset7

This is a dataset for a fold prediction task, similar to dataset 4. However, here the classification is only to seven possible folds. These data were previously assembled (Andreeva et al. 2020) and include 14,112, 1,568, and 3,921 training, validation, and test pairs, in which each pair includes a sequence and its associated fold. These data were taken from ProteinBERT (Brandes et al. 2022). As these data did not contain a validation set, we randomly sampled 10% of the training data to serve as a validation data.

#### Dataset8

Neuropeptides are peptides that are used for communication between neural cells and their peripheral cells (Burbach 2010). The vast majority of neuropeptides are translated as precursor neuropeptides and undergo cleavage and maturation events. Ofer and Linial (2014) have previously assembled a database of precursor neuropeptides and non-precursor neuropeptides from various animal species and developed a binary classification algorithm for predicting precursor neuropeptides given a set of protein sequences. We re-analyzed their data, which included 2,727, 303, and 337 training, validation, and test pairs, in which each pair includes a sequence and whether or not it is a neuropeptide.

Of note, datasets 4, 5, and 6 were previously analyzed by Rao et al. (2019) and Brandes et al. (2022) and datasets 7 and 8 were previously analyzed by Brandes et al. (2022).

### Tokenizer Implementation

We compared five different tokenizers: BPE, WordPiece and Unigram (Sennrich, Haddow, and Birch 2016; Schuster and Nakajima 2012; Kudo 2018) as well as the “words” and “pairs”. The three first tokenizers can be trained for specific data, i.e., they are data-driven. These were trained (on the training data) with default parameters (e.g., parameters that control the trimming of the Unigram program). The output of this stage is a dictionary for each transformer and dataset. All datasets were next encoded using the obtained dictionaries. This step was achieved using the HuggingFace library (Wolf et al. 2020). For each tokenizer, we evaluate the following dictionary sizes: 100, 200, 400, 800, 1,600 and 3,200. The output of each tokenizer and sequence is a vector of tokens (represented as integer numbers). Different sequences are represented by a different number of tokens, and in order for all vectors to be of the same length (a requirement of the transformers), a maximum size was set. Specifically, for datasets 1 the maximum size was set to 100 tokens. Similarly, for datasets 2, the maximum size was set to 512 tokens, for datasets 3 to 1,024 tokens and for datasets 4-8, to 512 tokens. For all classification and regression tasks, proteins longer than this size were trimmed and proteins shorter than this size were padded with a special token. These encoded data were next used to train transformers (on the training data) for the specific computational task associated with each dataset.

### Training the Transformers

Two different transformer architectures were considered for all classification and regression tasks: BERT (Devlin et al. 2019) and GPT (Radford et al., 2018). As the former resulted in higher performance on the validation data, we only present results obtained with BERT. In each case, the models are randomly initialized and trained on the tokenized training data of each dataset. Using the validation data, we have optimized several hyperparameters for each computational task: the number of layers, the number of attention heads, and the size of the hidden vector. The best performing configuration was with two hidden layers and two attention heads for all datasets. The size of the hidden vector was 128 for all datasets, except dataset 1, for which the optimal performance was with a vector of size 64. Transformer training and testing were implemented using the HuggingFace library (Wolf et al. 2020).

For dataset 3, which is associated with a sequence-to-sequence task, following Dotan et al. (2023), we relied on the “vaswani_wmt_en_de_big” architecture, with 6 hidden layers, 16 attention heads, and a hidden vector size of 1,024. The training was conducted with the Fairseq library (Ott et al. 2019). The learning rate, warmup values, and max tokens were set to: 5 × 10^−5^, 3,000, and 4,096, respectively.

### Comparison with Previous Models

We compared the performance of the different trained models with those obtained in previous studies. Specifically, datasets 4, 5, and 6 were each previously analyzed by applying three different models: ProteinBERT (Brandes et al. 2022), the Tasks Assessing Protein Embeddings (TAPE) transformer (Rao et al. 2019), and a biological model that relied on the LSTM architecture (Rao et al. 2019). Datasets 7 and 8 were previously analyzed using ProteinBERT. These previous works all used the “words” tokenizer. As we did not pre-train our models, for a fair comparison, the performance of these previous models was evaluated without pre-training.

### Evaluating the Performance on the Different Tasks

For classification tasks, we report Accuracy (ACC), Area Under the Curve (AUC), and Matthew’s Correlation Coefficient (MCC) (Matthews 1975). The latter is more suitable for unbalanced datasets as it considers the number of samples from each class:

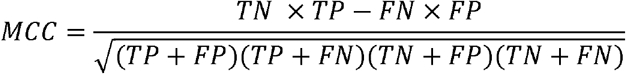

TN is the number of true negatives, TP true positives, FN false negatives and FP for false positives. We note that the range of MCC is from -1 to 1 while ACC and AUC range between 0 and 1.

For regressions tasks, we report the Spearman rank correlation, ranging from -1 to 1, where 1 reflects a perfect score.

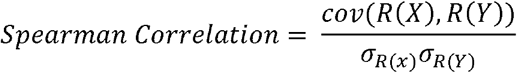

*X* and *Y* are the predictions and the labels, respectively. *R*(*z*) are the ranks of *Z, σ_z_* is the standard deviations of the *Z*, and cov(*Z, W*) is the covariances of the *Z* and *W*.

For the alignment task, we report the performance of the aligners with the Column Score (CS). The CS is measured by counting the number of columns in the inferred alignment that have a matching column in the true alignment, out of the total number of columns in the true alignment (Penn et al. 2010). The range of the CS is between zero and one. We also report the coverage score:

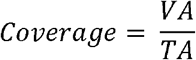

where VA is the number of valid alignments and TA is the total number of alignments tested. When using transformers to align biological sequences, it is possible that the transformer erroneously creates invalid alignments. For example, if the input sequences are “AAG” and “AAC”, the transformer may output a pairwise alignment in which “AA–G” is aligned to “AAC”. This is clearly an invalid alignment as the number of characters in all alignment rows should be identical. In addition, each alignment row should be identical to the original corresponding (unaligned) sequence after removing all of it gaps. In rare cases, this is not the case, and these alignments are also considered invalid (Dotan et al. 2023). Of note, all alignments in the training data are valid alignment. Thus, higher coverage suggests better learning from the training data.

### Visualizing the Signals Within Protein Superfamilies

For the task of classifying sequences to superfamilies, the trained transformer allows highlighting the positions and signatures (tokens) that contribute most for distinguishing one superfamily from the others. We used the Captum library (Kokhlikyan et al. 2020), which allows interpreting trained transformers regarding their decision making. To this end, the integrated gradients method (Sundararajan, Taly, and Yan 2017) was used to calculate the importance of each of the input tokens (we used the default number of steps which is 50). For example, in the context of superfamily classification, how important each token is for the correct identification of a specific superfamily. Unlike standard motifs used in computational biology, here the algorithm can highlight both tokens whose inclusion in the protein sequence supports a classification to a specific superfamily and tokens whose exclusion supports the classification. To do this, we first identified the positions of tokens with high attribution scores in each sequence (absolute value above 0.2). Then, we created histograms for each family to see where these high score tokens were located. We next searched for specific amino acids patterns by using a sliding window approach. Moving along the sequences a window of size 15 amino acids, we searched for the presence of a specific token in at least eight out of 100 sequences within each superfamily. If so, we added a label for the token at that location. This information can be used to identify the locations of signals on protein sequences. The transformer used was the best performing one, trained on the BPE-tokenized data with 1,600 vocabulary items.

### Training on the BFD Dataset

The BFD dataset is currently the largest public dataset of protein sequences, comprising over 2.2 billion sequences and 400 billion amino acids (Steinegger and Söding 2018). This dataset includes multiple sources of sequences that were aligned to longer sequences using MMseqs2 (Steinegger and Söding 2017) and filtered based on sequence identity and number of sequences per cluster. After obtaining this dataset, we preprocessed the sequences by removing gap characters (“–”). This preprocessing phase resulted in a file of approximately 400 GB, containing pure proteins. We trained the BPE, WordPiece, and Unigram tokenizers on this dataset. Due to the large memory required for this task, we utilized the AMD EPYC 7H12 machines, with 256 cores and approximately 1 Tb RAM. Due to memory and run time limitation, we trained the different tokenizers on subsamples with increasing sizes, ranging from 1,000 to 10,000,000 sequences.

### Data Availability

Code, data and trained tokenizers are available on https://github.com/idotan286/BiologicalTokenizers.

## Results

### Effectors and Superfamilies Classifications

We first evaluated the performance of the various tokenization algorithms for the task of classifying proteins to those that are effectors and those that are not (dataset 1). Figure 2a shows the performance, as measured by the MCC score, for the various tokenizers. It also shows the reduction in the length of the encoded proteins. The optimal performance, with an MCC score of 0.62, was obtained using the WordPiece tokenizer with a dictionary size of 400 tokens. Compared with the default “words” tokenizer, it is both more accurate (the MCC of the “words” tokenizer was only 0.27), and it led to a 1.9-fold reduction in sequence length (i.e., the number of tokens). In fact, the “words” dictionary had the lowest length reduction (as expected) and the second lowest MCC score. The highest fold reduction of 2.4 in length was obtained with the WordPiece tokenizer when using 3,200 tokens, albeit with a reduction of 0.12 in the MCC score compared to the best tokenizer. We note that this tokenizer had higher performance and higher fold reduction compared to the “pairs” tokenizer, which in turn was better than the “words” tokenizer (Figure 2a).

**Figure 2:**
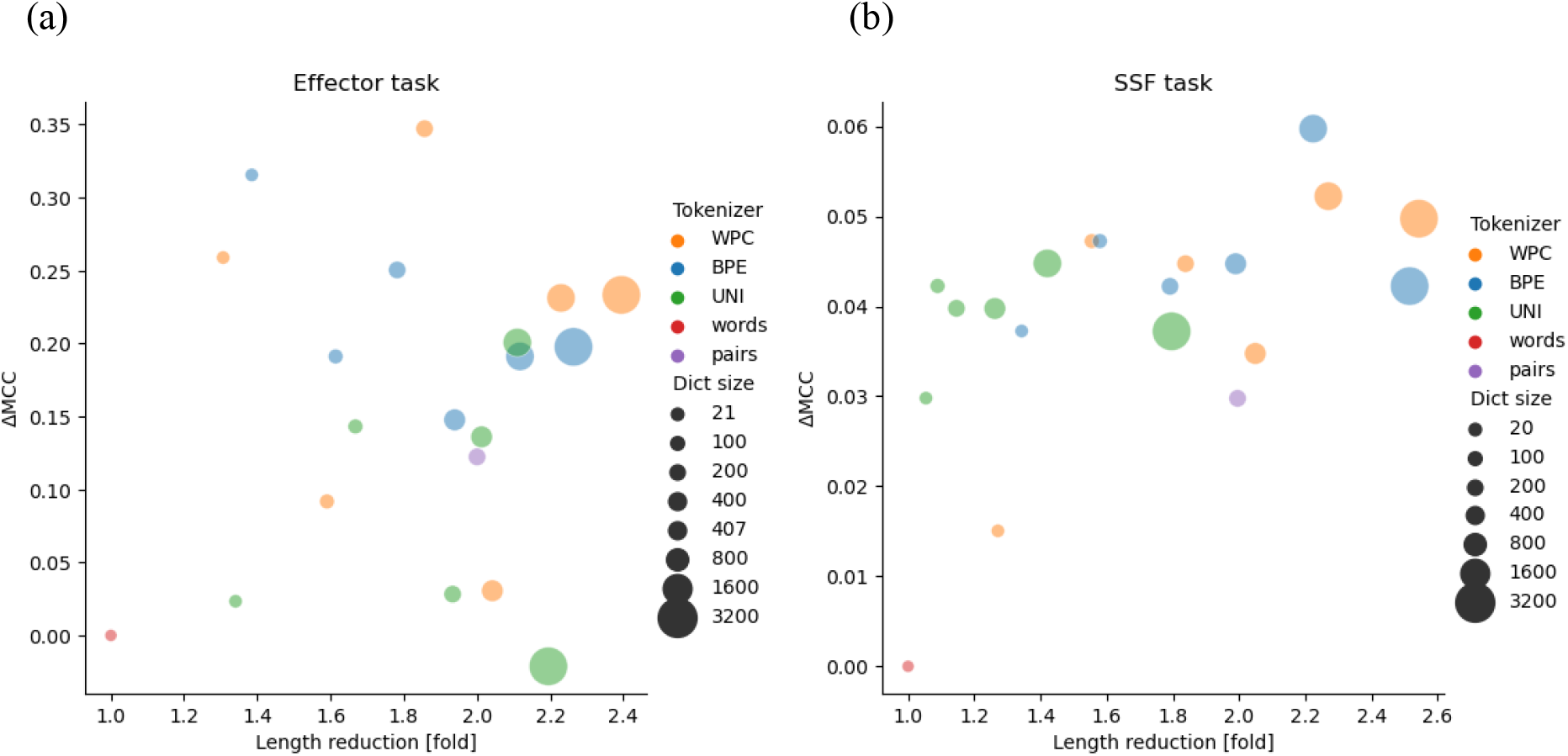
Panels (a) and (b) show the results of the trained transformers on the effector and superfamilies (SSF) classification tasks, respectively. A different color is assigned for each different tokenizer: BPE, WordPiece (WPC), Unigram (UNI) and the baseline tokenizers: “words” and “pairs”. The dictionary size (Dict size) is demonstrated by the size of the circle. The x axis indicates the fold-reduction in number of tokens used (i.e., the “length”) relative to the “words” tokenizer. The y axis indicates the improvement in MCC score relative to the “words” tokenizer. The “word” tokenizer had MCC scores of 0.27 and 0.935 for the effector and SSF tasks, respectively. The closer the dictionary is to the right upper corner, the better it is, as it has higher length reduction and higher performance.

Figure 2b demonstrates the performance of applying the different tokenizers on the multiclass classification (dataset 2). The BPE tokenizer resulted in lower sequence length compared to Unigram. While the 3,200 tokens Unigram dictionary led to only a 1.79-fold reduction in sequence length, BPE that has the same dictionary size, led to over 2.5 folds reduction in sequence length. The best performing transformer used the BPE tokenizer. It was trained on a dictionary containing 1,600 tokens and had a very high performance, with an MCC score of 0.995. It led to a 2.2-fold reduction in the number tokens. Of note, the transformer trained with the “words” tokenizer has lower performance compared to transformers trained with alternative tokenizers. Similarly, most transformers using data-driven tokenizers outperformed the “pairs”-based dictionary. Interestingly, the dictionaries of 100 and 200 tokens of BPE had a resulted in higher length-reduction compared to the same size dictionaries of WordPiece. This was reverted when comparing larger dictionaries with 400 or more tokens.

### Alignment

Next, we evaluated the impact of tokenizing nucleotide sequences on alignment accuracy and coverage (see Methods). Figure 3a shows the performance of the transformers on the alignment dataset. The accuracy of all transformers (measured by the CS score) were very high (above 0.98), and the differences among the different transformers were very small (∼0.007 difference in the CS score between the best and the worst transformer). These high scores suggest that all transformers could reliably aligned the analyzed sequences.

**Figure 3:**
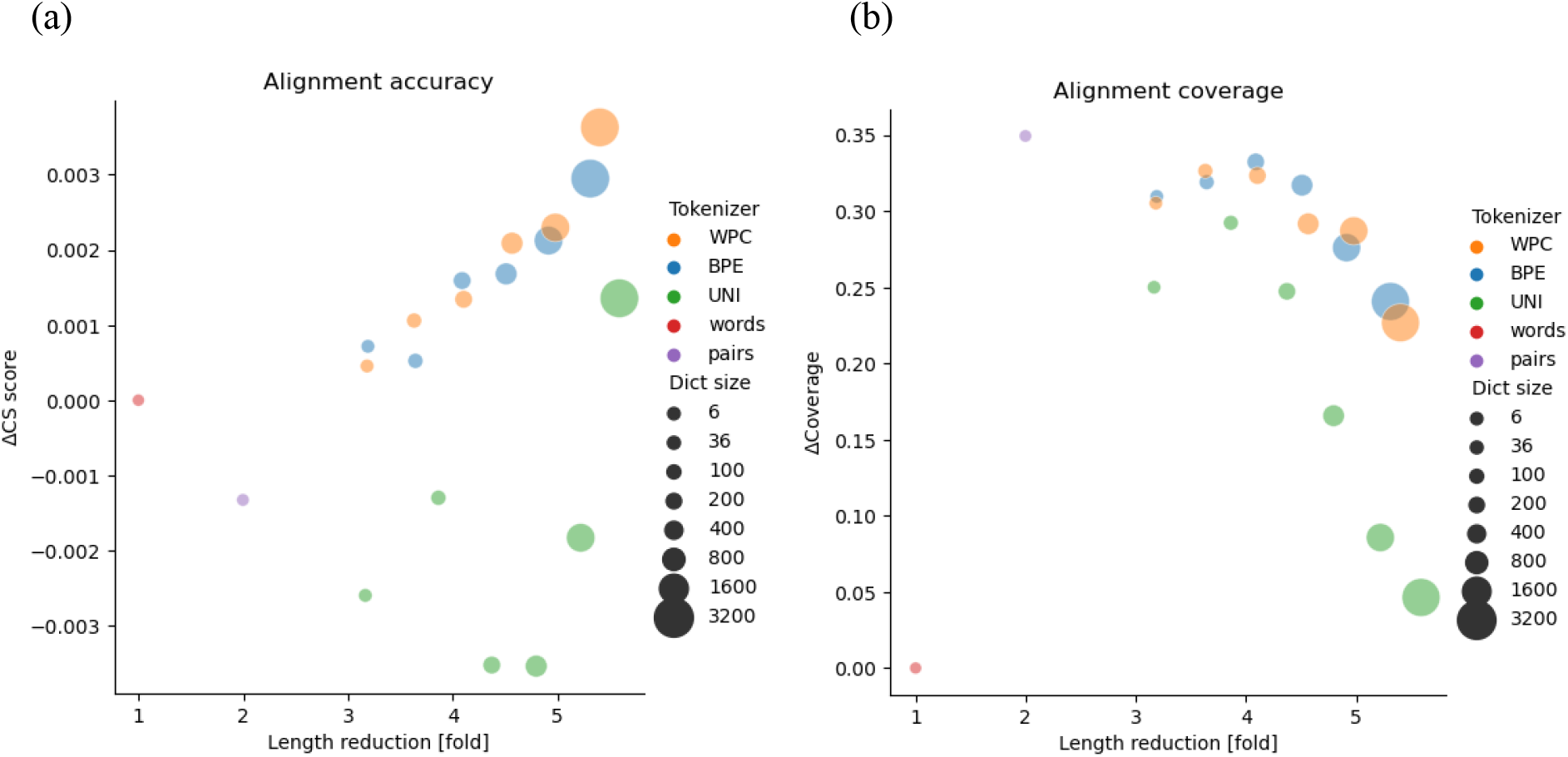
Results of the transformers trained on the alignment dataset preprocessed by different tokenizers: BPE, WordPiece (WPC), Unigram (UNI) and the baseline approaches: “words” and “pairs”. Panels (a) and (b) report the performance as measured by the CS score and the coverage, respectively. The “word” tokenizer had CS score of 0.99 (panel a), and a coverage of 0.59 (panel b). Of note, the CS score is only computed on valid alignments.

Figure 3b shows the coverage of the transformers on the alignment dataset. The coverage is calculated as the number of valid alignments divided by the number of tested alignments (see Methods). Large differences in coverage were observed between the worst and best transformers: the transformer using the “words” tokenizer obtained a coverage of 0.59, while the “pairs” had the highest coverage of 0.941. The best performing transformer using a data-driven tokenizer was the BPE transformer with a dictionary size of 400, resulting in a coverage of 0.924. However, this BPE tokenizer reduced the number of tokens by more than four, while the baseline reduced the number of tokens by only two. Thus, the BPE with 400 tokens had only half as many tokens as the “pairs” baseline. As can be seen from the figure, there is a trade-off between reduction and performance. Careful examination of the BPE, WordPiece, and Unigram tokenizers shows that increasing the vocabulary size increases the coverage and reduction fold, but once the size reaches a few hundred (400, 200, 200 for BPE, WordPiece and Unigram, respectively), the coverage begins to decrease. Even dictionaries of 100 or 200 tokens have large impact on the length of the encoded sequences, slightly more than three-fold.

### Comparison of Different Models on additional Classification and Regression Tasks

Figure 4a illustrates the results gained on the remote homology classification (dataset 4) (Rao et al. 2019; Hou, Adhikari, and Cheng 2018). The results of the regression task of predicting the log-fluorescence of proteins (dataset 5) are demonstrated in Figure 4b (Rao et al. 2019; Sarkisyan et al. 2016). Figure 4c shows the results of the proteins stability regression task (dataset 6) (Rao et al. 2019; Rocklin et al. 2017). The results of training the transformers on the tokenized data were compared to the previously published results of ProteinBERT (Brandes et al. 2022), TAPE (Rao et al. 2019), and LSTM (Rao et al. 2019) without their pretraining (Figure 4). Comparing the TAPE transformer with one of a transformer that used a data-driven tokenizer, specifically WordPiece with the 400 tokens, revealed that the later was both more accurate and used less tokens in all three computational tasks (Figure 4). Of note, TAPE includes seven times more free parameters than the transformer using WordPiece. Carefully examining the performance of ProteinBERT (Brandes et al. 2022) reveals it has the best performance on the fluorescence (Figure 4b) and stability (Figure 4c) tasks and the lowest performance on the remote homology task. Of note, while the ProteinBERT, LSTM and TAPE have 16 million (M), 38M and 38M parameters, respectively, the remaining transformers studied have only 5M free parameters. One of the main advantages of using the optimized tokenizers was demonstrated in the fluorescence task (Figure 4b), where using dictionaries with a small number of tokens (such as 3,200 for BPE and WordPiece) was able to significantly reduce the length of the input sequences (by up to 20 fold) compared to the tokenizers used in ProteinBERT, LSTM, and TAPE.

**Figure 4:**
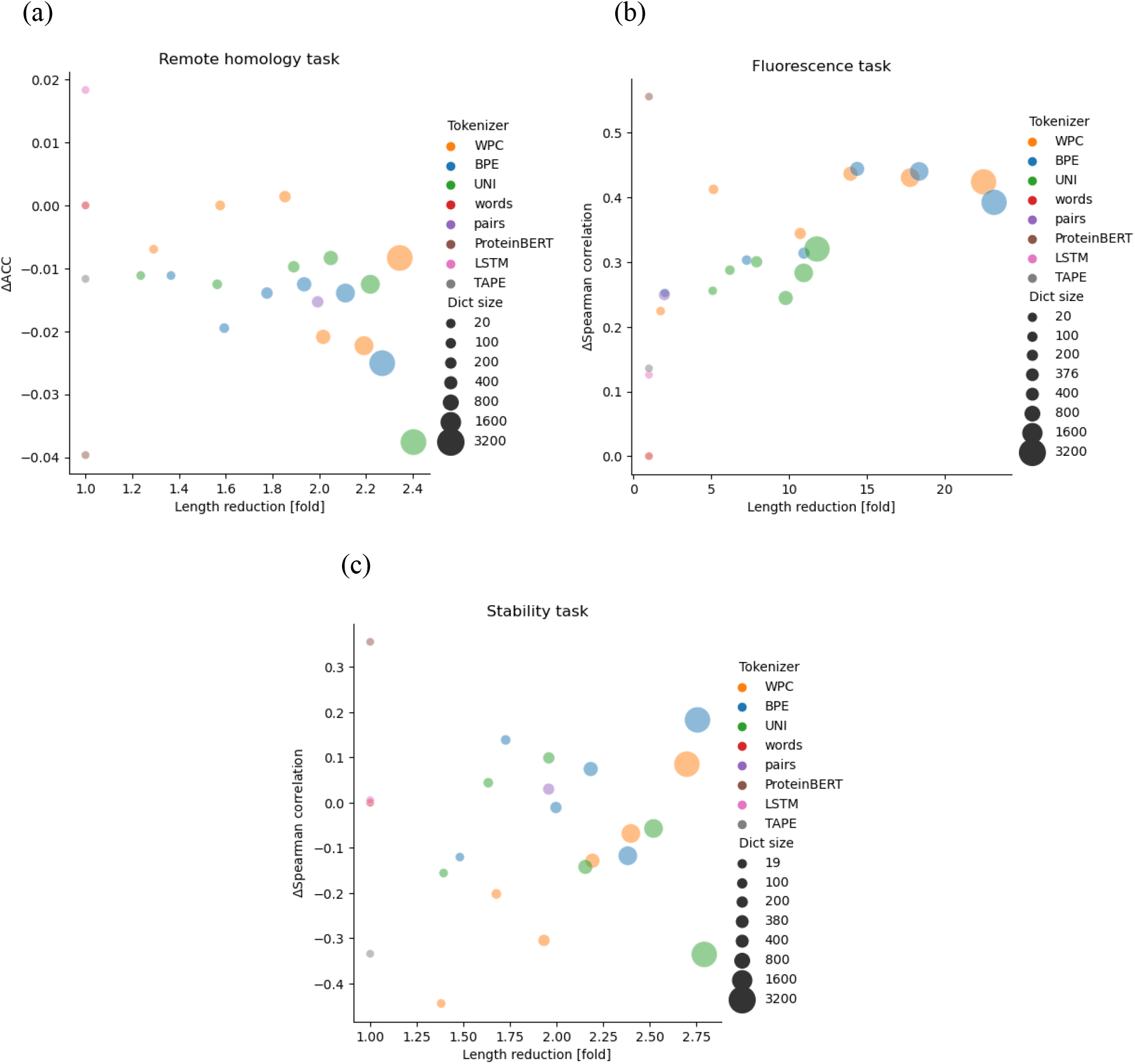
Evaluating the performance of the different tokenizers and comparing them to the previously tested models: ProteinBERT, LSTM and TAPE. The x-axis is the length reduction, and the y-axis is the performance of the transformer trained on the same dataset. Panels (a), (b), and (c) display the results of datasets 4, 5, and 6, respectively. The “words” tokenizer had an accuracy (ACC) score of 0.101, spearman correlation scores of 0.084 and 0.274 for the remote homology task, fluorescence task and the stability task, respectively.

We compared the various tokenization techniques on two additional tasks, previously analyzed in the ProteinBERT study (Brandes et al. 2022): fold prediction and neuropeptide classification (Figure 5). For the fold prediction task (Figure 5a), we observe a trade-off between the performance and the dictionary size. Of note, the transformer with the Unigram tokenizer (with 100 tokens) was both more accurate and used less tokens than both “words” and ProteinBERT. For the neuropeptide prediction task, the ProteinBERT transformer performed worst (Figure 5b) than all other transformers. Here is a single transformer (the WordPiece with 3,200 tokens) was both more accurate and had the highest length reduction compared to all other methods.

**Figure 5:**
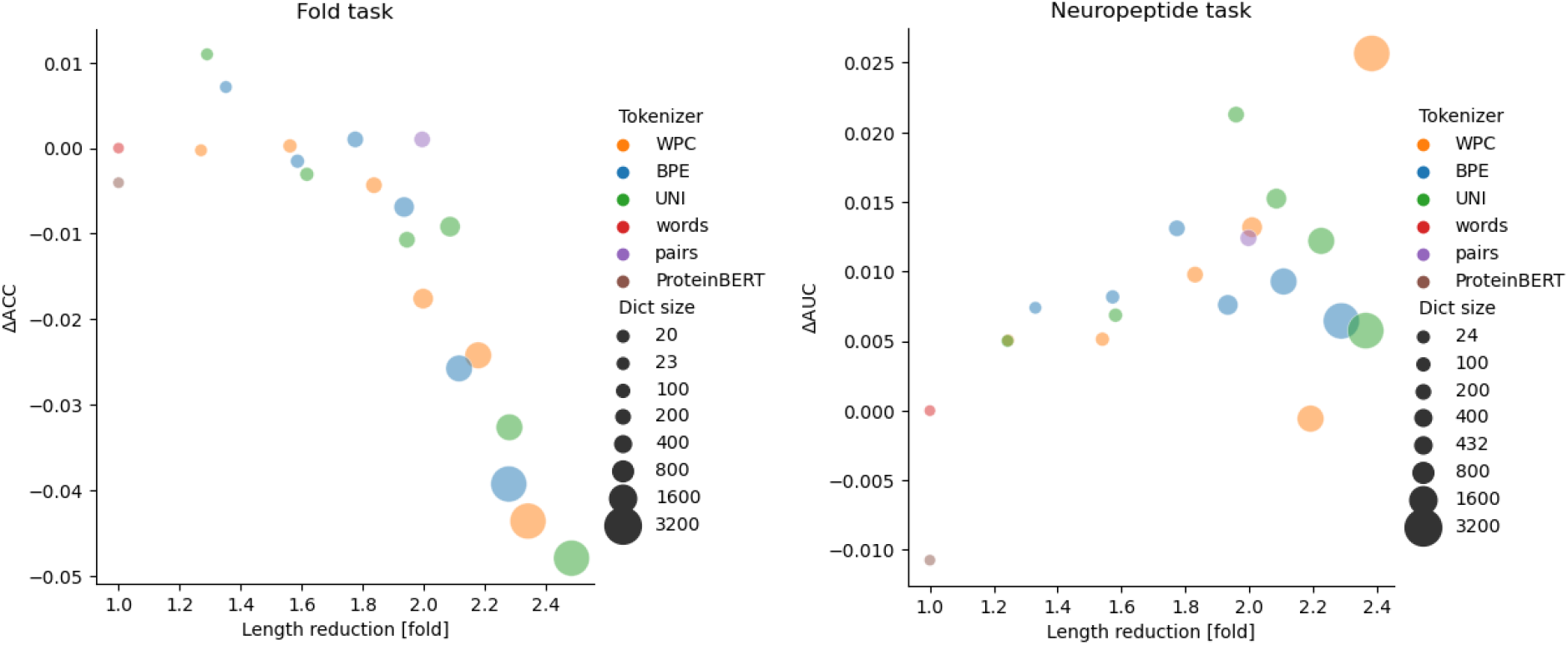
We evaluated the tokenization on two datasets proposed in ProteinBERT: fold structure classification and neuropeptide identification. The x-axis and the y-axis refer to the length reduction and the performance, respectively. The “words” tokenizer had an ACC score of 0.59, and an AUC score of 0.95 for the fold task, neuropeptide task, respectively.

Our results suggest that for some datasets a tradeoff between accuracy and fold reduction exists, while for some tasks, using data-driven tokenizers can be beneficial in both aspects (accuracy and length reduction). In addition, our results suggest that the ProteinBERT architecture may be more suited for regression tasks, than to other tasks such as classification.

### Identification of Contributing Signals and Their Visualization

One of the key disadvantages of using transformers is that they are highly non-linear and difficult to interpret. Biological sequences contain signals in specific locations that are important for determining their structure and function. Identifying these signals remains a challenge. Better understanding of these signals should result in a better mapping between a protein sequence and its structure and function, thus contributing to protein function prediction, classification, and design of novel proteins. By training a transformer to predict specific classes, we could apply interpretation tools to identify those signals. Figure 6 displays the resulting interpretation of each superfamily, specifically enrichment and depletion of specific tokens as a function of the protein length. For example, a “PKK” at the N terminus of a protein suggests it belong to the superfamily SSF101152. One of the key differences from the signatures that appear in Pfam (Mistry et al. 2021) is that here we do not rely on a multiple sequence alignment, thus accounting for variability in the position of specific tokens along the length of the protein. In addition, dependencies among tokens are accounted for. Finally, depleted tokens can be identified, e.g., the presence of the token “ER” at the N terminus of proteins suggests it is not SSF100879.

**Figure 6:**
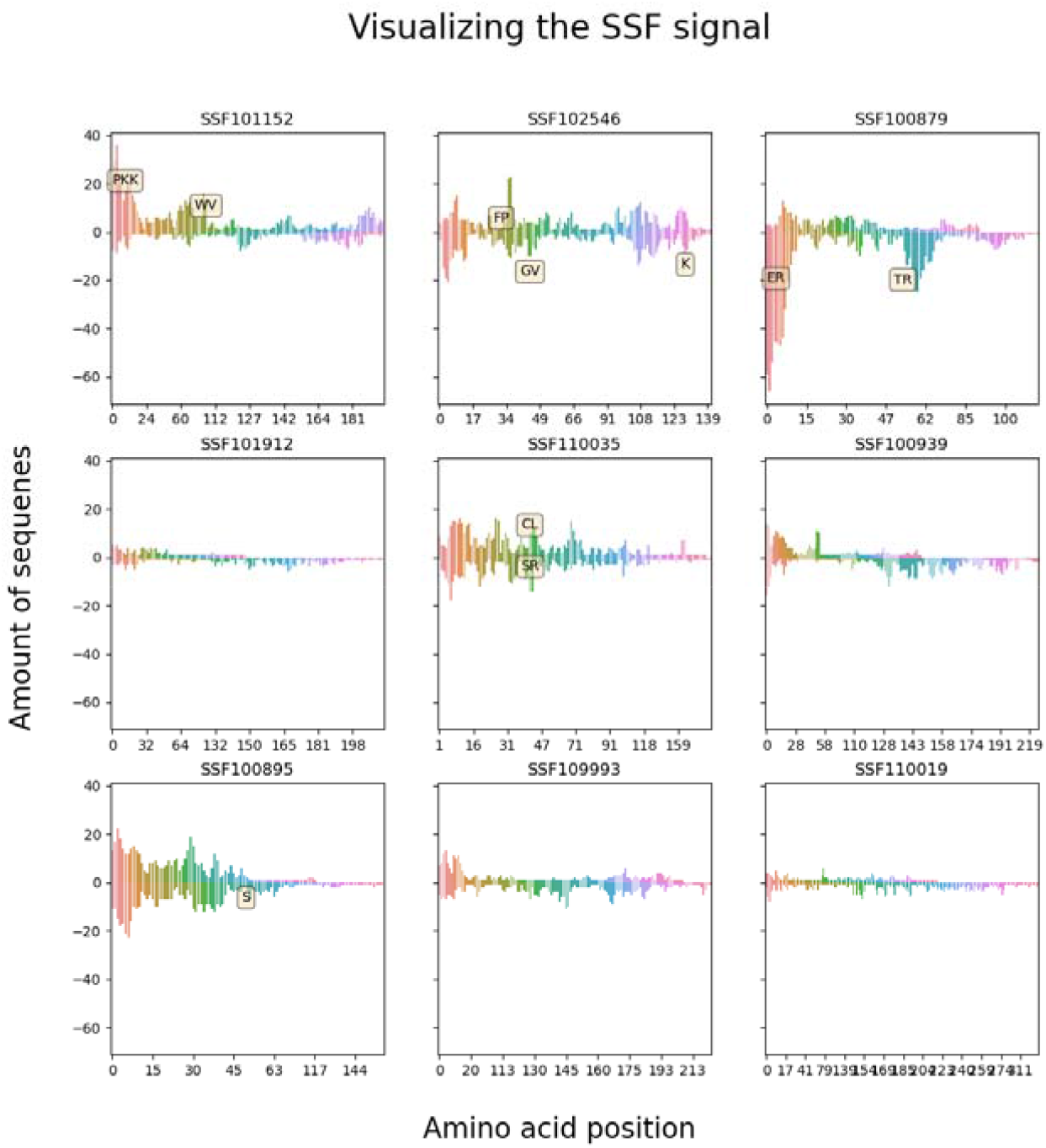
visualizing the importance features of each of the superfamilies. Each of the nine graphs correspond to different family. In each graph, the x axis is the amino acid position, and the y axis the number of sequences that had an important feature in this position. We added labels with specific tokens if they are repeated in several different sequences. The graphs were created by 100 test protein sequences for each superfamily.

### Tokenizing the BFD Dataset

In our previous experiments, we showed the effect of tokenizing the input on specific tasks, i.e., for each task we trained data-specific tokenizers. Here, we aimed to train tokenizers on a very large dataset, which may be important in cases where there are not enough sequences for a specific task, or as input for the next generation of biological pre-trained models. To this end, we trained BPE, WordPiece and Unigram tokenizers on samples of proteins from the 2.2 billion protein sequences of the BFD dataset (Steinegger and Söding 2018). We evaluate the average sequences length as a function of the vocabulary size and number of sequences in the training data (Figure 7). Increasing the size of the vocabulary resulted in a sharp decrease in the average numbers of tokens per protein, thus enabling processing longer biological sequences with the similar memory requirement. In addition, the increase of the training data resulted in smaller values of tokens per protein. The average number of tokens per protein converged after training on 1,000,000 samples. When using the largest dictionaries (51,200 tokens), the BPE, WordPiece and Unigram reduced the average length by 15.6%, 16.8 and 14.9%, respectively. Among all tokenizers, Unigram was most influenced by the training and vocabulary sizes: it has the highest number of tokens per proteins (182.4) when using small training size (1,000 sequences) and 100 tokens in the vocabulary). Yet, when the training data was 10^7^ sequences and the dictionary size higher than 50,000 tokens, it obtained the best average length (53.1 tokens for protein).

**Figure 7:**
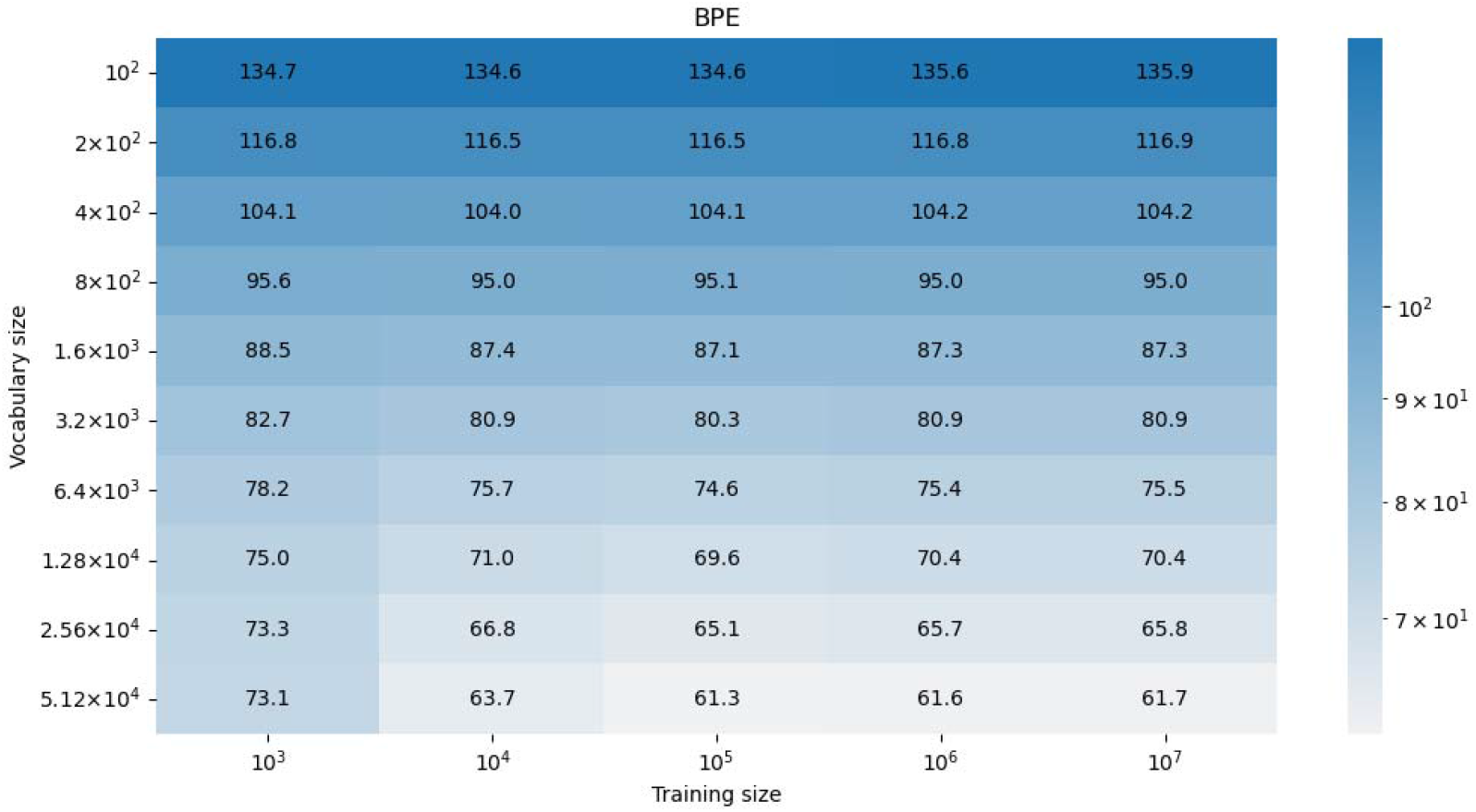
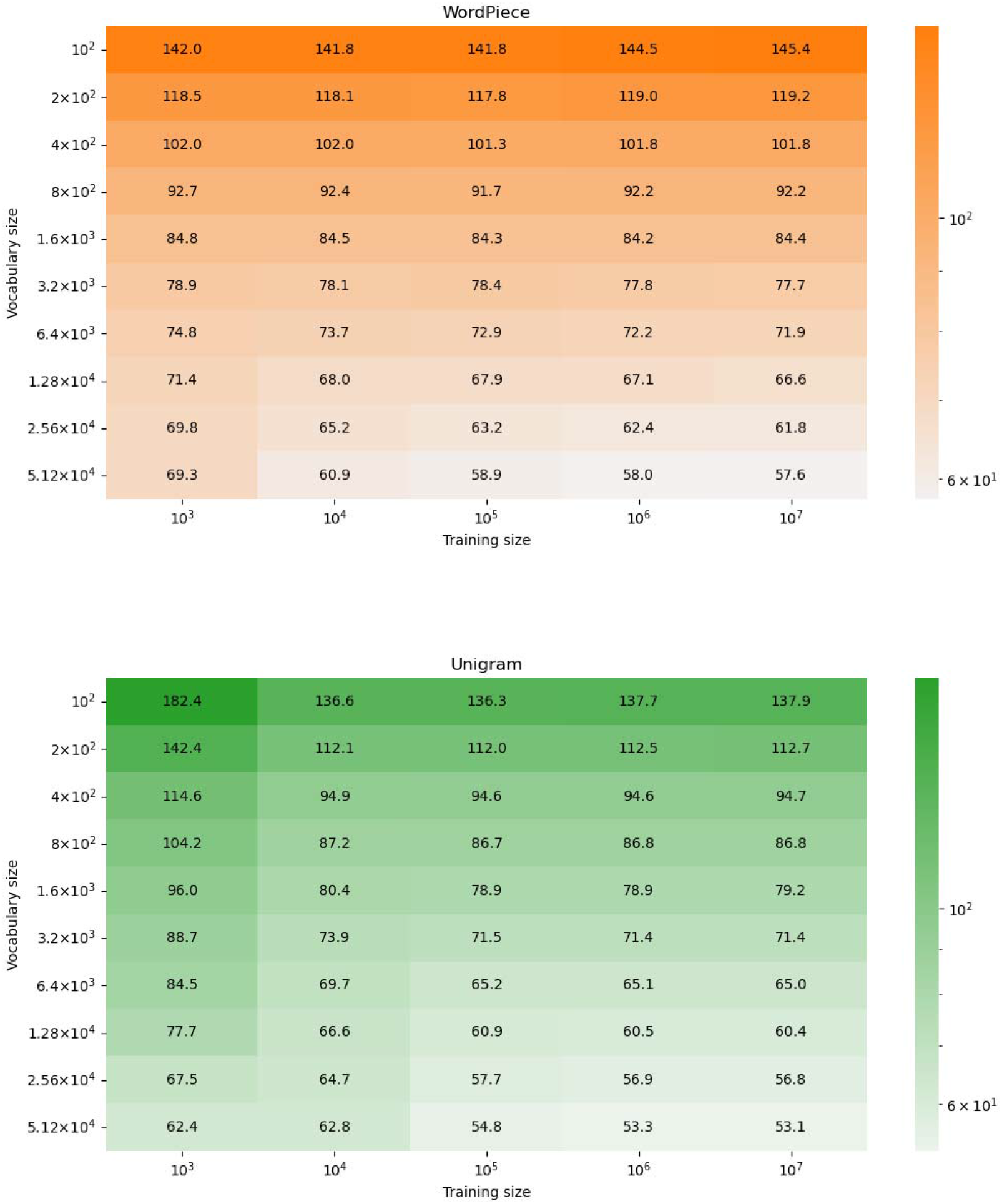
Effect of vocabulary size and number of training samples on the three tokenizers: BPE, WordPiece and Unigram. The darker the color the higher the average number of tokens per protein. Increasing the vocabulary and the training size reduces the number of tokens per protein for all of the tested tokenizers.

### Comparing Specific versus General Data-Driven Tokenizers

Above we trained two types of data-driven tokenizers. The first type, which we term “specific”, was trained on small datasets (e.g., effectors), while the second type, which we term “general” was trained on very large number of protein sequences, not related to a specific computational task. We next aimed to determine whether it is beneficial, for specific tasks, to use general versus specific data-driven tokenizers. To this end, we compared the performance between the specific and the general versions of the trained transformers BPE, Unigram, and WordPiece on seven computational tasks (the alignment task is based on nucleotide rather than protein sequences and was hence not included here). For all tasks, the specific type outperformed the general type, and the results were statistically significant for six out of the seven tasks (Figure 8). For some tasks such as the remote homology task, using the general tokenizers resulted in very poor performance compared to the specific tokenizers. We hypothesize that tasks involving proteins with similar domains would be more affected by using task-specific tokenizers, while tasks encompassing proteins across the tree of life would likely show similar results to those using the BFD-trained tokenizers..

**Figure 8:**
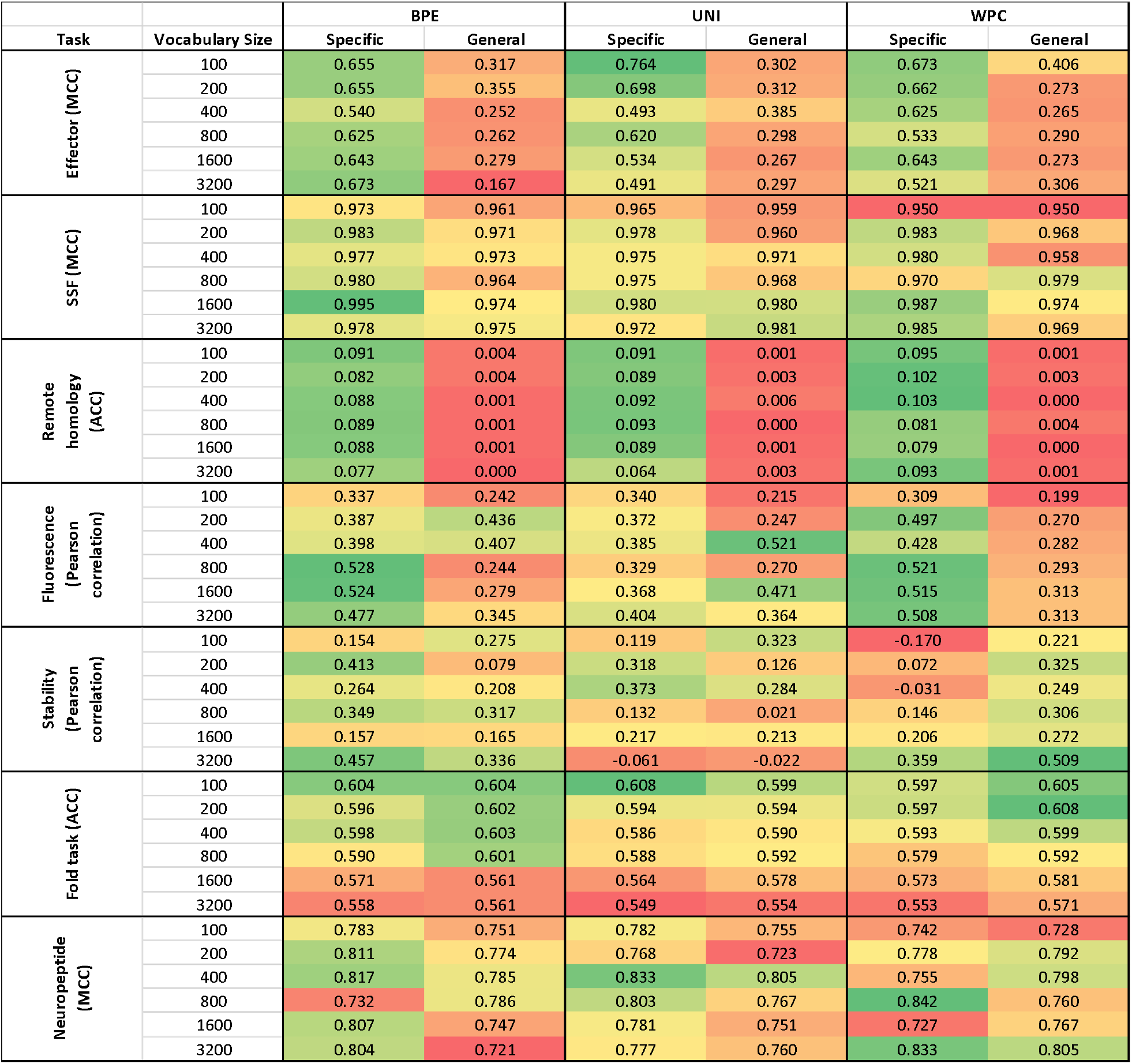
Comparing the performance of transformer trained on data encoded by specific trained tokenizers (“Specific”) and tokenizers trained on the BFD dataset (“General”). The evaluation was conducted on seven datasets, utilizing three tokenizer types: BPE, Unigram, and WordPiece. For each tokenizer, multiple vocabulary sizes were tested: 100, 200, 400, 800, 1,600, and 3,200. The performance is represented by the green and red colors, where a higher intensity of green indicates better performance. Each task is individually colored to facilitate comparison. To quantify the differences between the general and specific tokenizers, we performed paired t-tests and obtained the following p-values: 8.23^-11^, 0.001, 5.07^-18^, 0.0016, 0.356, 0.0004, 0.025, for datasets 1, 2, 4, 5, 6, 7, and 8, respectively

## Discussion

Our results clearly indicate that the use of tokenization in the analysis of biological datasets can improve performance. This was demonstrated for all tested datasets and for all types of analyses (classification, regression, and sequence-to-sequence). However, no single tokenization method was optimal for all datasets, emphasizing the need to evaluate different tokenizers for each data and learning task.

Transformers often cannot analyze sequences above a specific threshold length. It is common to segment longer sequences to subsequences shorter than this threshold, thus bypassing this restriction. However, this fragmentation prohibits the model from analyze the entire input data, and can thus potentially decrease performance. Our results show that fragmentation can be avoided by tokenizing the data, i.e., tokenization allows architectures to expend their capacity to substantially longer proteins and DNA sequences, as was recently shown in DNABERT-2 (Zhou et al. 2023).

In this study we also demonstrated how important biological information can be extracted for post-analysis of trained models applied to specific learning tasks. For example, we could detect specific signature for protein super-family classification. One of the benefits of the proposed approach compared to motifs in the form of Profile Hidden Markov Models is that it does not rely on a multiple sequence alignment, which may be unreliable, especially when highly diverged sequences are analyzed.

In this study, we examined the impact of tokenization at the molecular level. We hypothesize that tokenization has the potential to be applied to various forms of discrete biological data, such as genes (Miller, Stern, and Burstein 2022). Additionally, the incorporation of character compression into classical algorithms used in biology, such as Blast (Altschul et al. 1990) and Kraken-2 (Wood, Lu, and Langmead 2019), should be considered in order to decrease running times.

This work represents the initial phase of studying how tokenization impacts biological language models. Our study demonstrates that data-driven tokenizers should be considered, both for accuracy and for length reduction. Our work also shows that there is no single data-driven tokenizer that outperformed all the others. We demonstrate that the effect of tokenizing the sequence depends on the specific task, the data type and size, and the tokenization algorithm applied. In future work, it would be interesting to compare Large Biological Models (LBMs) performance which were pretrained with various tokenization algorithms, i.e., we speculate that in the future there will be several alternative LBMs, each pretrained with a different tokenization algorithm, and users can test which LBM is best suited to their computational task. Our study further suggests that future studies comparing the performance of new emerging transformer architectures on biological data, should include different tokenizers as a critical component in their evaluation.

## Acknowledgments

T.P. and Y.B. have received funding from the Israel Science Foundation (Grants 2818/21 and 448/20, respectively). Y.B. was supported by an Azrieli Foundation Early Career Faculty Fellowship. E.D. was supported in part by a fellowship from the Edmond J. Safra Center for Bioinformatics at Tel Aviv University.

